# The P2 protein of wheat yellow mosaic virus acts as a viral suppressor of RNA silencing to facilitate virus infection in wheat plants

**DOI:** 10.1101/2023.12.07.570682

**Authors:** Dao Chen, Hui-Ying Zhang, Shu-Ming Hu, Zheng He, Yong Qi Wu, Zong-Ying Zhang, Ying Wang, Cheng-Gui Han

## Abstract

Wheat yellow mosaic virus (WYMV) causes severe viral wheat disease in Asia. The WYMV P1 protein encoded by RNA2 has viral suppressor of RNA silencing (VSR) activity to facilitate virus infection; however, VSR activity has not been identified for P2 protein encoded by RNA2. In this study, P2 protein exhibited strong VSR activity in *Nicotiana benthamiana* at the four-leaf stage, and point mutants P70A and G230A lost VSR activity. Protein P2 interacted with calmodulin (CaM) protein, a gene-silencing associated protein, while point mutants P70A and G230A did not interact with it. Competitive bimolecular fluorescence complementation and competitive co-immunoprecipitation experiments showed that P2 interfered with the interaction between CaM and calmodulin-binding transcription activator 3 (CAMTA3), but the point mutants P70A and G230A could not. Mechanical inoculation of wheat with *in vitro* transcripts of WYMV infectious cDNA clone further confirmed that VSR-deficient mutants P70A and G230A decreased WYMV infection in wheat plants compared with the wild type. In addition, RNA silencing, temperature, and autophagy had significant effects on accumulation of P2 protein in *N. benthamiana* leaves. In conclusion, WYMV P2 plays a VSR role in wheat and promotes virus infection by interfering with calmodulin-related antiviral RNAi defense.

**One-sentence summary:** WYMV P2 protein exerts VSR activity by interfering with the CaM–CAMTA3 interaction to facilitate virus efficient systemic infection in wheat plants.

## Introduction

Wheat is one of the main grain crops in Asia, and the wheat disease caused by wheat yellow mosaic virus (WYMV) can significantly reduce wheat yield (Chen and Ruan, 1989; Kanyuka et al., 2003; Kühne, 2009; Miller et al., 1992; Slykhuis, 1970; Usugi et al., 1979). The symptoms of WYMV disease include yellowing, chlorosis, dwarfing, and even death in severe cases. When the temperature rises to about 10 ℃ in February every year in Asia, the WYMV disease enters its peak season (Chen et al., 1999; Lei and Chen, 1998; Namba et al., 1998; Yu et al., 1995; Yu et al., 1999). WYMV belongs to the genus *Bymovirus* in the family *Potyviridae*; the genus includes six viruses – WYMV, wheat spindle streak mosaic virus (WSSMV), barley mild mosaic virus (BaMMV), barley yellow mosaic virus (BaYMV), oat mosaic virus (OMV), and rice necrosis mosaic virus (RNMV) – which are all spread in the field by *Polymyxa graminis*. Among them, WYMV and WSSMV only infect wheat (Chen and Chen, 2002; Han et al., 2000; Jiang and Kan et al., 2020; Kanyuka et al., 2003).

WYMV is a curved linear virus particle consisting of two single-stranded sense RNAs; the complete genomic sequences of four WYMV isolates from China and Japan have been reported (Chen et al., 1999; Namba et al., 1998; Ohki et al., 2019; Yu et al., 1999; Zhang et al., 2021). The WYMV RNA1 has an open reading frame and encodes a polymeric protein composed of 2404 amino acids; the polyprotein is digested by enzymes to produce 10 mature proteins, including P3, P3N-PIPO, PIPO, 7K, CI, 14K, VPg, NIa, Nib, and CP (Chen et al., 1999; Namba et al., 1998; Yu et al., 1999). Protein P3 is associated with virus infection (Chung et al., 2008; Luan et al., 2019; Yu et al., 2021); 7K is involved in WYMV replication (Rodriguez Cerezo et al., 1993; Yu et al., 2021); CI helps WYMV to form inclusion bodies in wheat cells (Ohki et al., 2019; Rodriguez Cerezo et al., 1997); 14K and NIa-Vpg contribute to WYMV replication (Li and Shirako, 2015; Schaad et al., 1997; Sun et al., 2013); NIb has the function of interfering with ABA signaling pathways (Zhang et al., 2019); and CP facilitates WYMV movement in wheat (Robaglia et al., 1989; Sun et al., 2013).

The WYMV RNA2 contains an open reading frame of 2712 nt, encoding a polyprotein of about 101 kDa with a cleavage site at amino acids (AA) 252–256, then the polyprotein is cleaved to produce P1 (28 kDa) and P2 (72 kDa) proteins (Namba et al., 1998; Oh and Carrington, 1989; Yu et al., 1999). Previous studies showed that WYMV P1 is a necessary factor for systemic infection of WYMV, and P1 may enhance RNA1 replication (You and Shirako, 2010). It was recently found that RNA silencing can be regulated by the signal cascade of calmodulin (CaM) protein binding with calmodulin-binding transcription activator 3 (CAMTA3) protein; the CaM– CAMTAs interaction upregulates transcription of RNA-silencing related genes such as those encoding dicer1 (DCL1), bifunctional nuclease 2 (BN2), Argonaute protein (AGO), and RNA-dependent RNA polymerase 6 (RDR6), which enhance the resistance of *Nicotiana benthamiana* plants (Wang et al., 2021; Wang et al., 2022). Our laboratory research showed that WYMV P1 protein also has viral suppressor of RNA silencing (VSR) activity, and plays a suppressor role by interfering with calmodulin-related antiviral RNAi defense, confirming that its VSR activity contributes to the systemic infection of WYMV in wheat (Chen et al., 2023). The P2 proteins of BaMMV, OMV, and BaYMV may be related to fungal transmission (Jacobi et al., 1995; Peerenboom et al., 1992; Zheng et al., 2002). Previous studies have shown that WYMV P2 is involved in WYMV replication and systemic movement (Li and Shirako, 2015; Sun et al., 2014; You and Shirako, 2010). The BaYMV P2 can promote viral systemic movement and symptom production in infected hosts (You and Shirako, 2010).

The amino acid sequences of P1 (WYMV) and P2 (WYMV) were compared with HC-Pro (TuMV), showing amino acid sequence identity of 9.8% and 9.23%, respectively (Supporting Figure 1). Recently, it was reported that areca palm necrotic spindle spot virus (ANSSV) of *Potyviridae* encodes a specific concatenated precursor cysteine protease (HCPro1–HCPro2); it was found that HCPro1 and HCPro2 differentially inhibit the antiviral immune response mediated by host RNA silencing and cooperatively maintain virus infectivity (Qin et al., 2020). Therefore, we speculate that WYMV (P1–P2) is similar to ANSSV (HCPro1–HCPro2), and P2 also has VSR activity. In this paper, Agrobacterium infiltration experiments showed that WYMV P2 protein had local and systemic VSR activities. Further studies showed that P2 interfered with the CaM–CAMTA3 interaction to exert VSR activity, and confirmed that its VSR activity was involved in the effective systemic infection of WYMV. The study results provide new insights and solutions for further understanding the role of P2 in WYMV infection.

## Results

### WYMV P2 protein has VSR activity

The proteins encoded by WYMV RNA1 and RNA2 are shown in Figure 1A. The VSR activity of WYMV P2 in *N. benthamiana* leaves at the four- and eight-leaf stages was observed under ultraviolet (UV) lamp. Empty vector (EV) was included in the experiment for comparison as a negative control. Compared with the negative control (EV+smGFP), which had almost no fluorescence, the experimental group (P2+smGFP) had light yellow fluorescence. However, fluorescence of (P2+smGFP) was more obvious in the four-than the eight-leaf stage (Figure 1B). Western blotting showed that P2 and GFP expression levels in the (P2+smGFP) group were significantly higher at the four-than at the eight-leaf stage (Figure 1C). Expression of P2 protein was lower in older (eight-leaf stage) than in younger *N. benthamiana* (four-leaf stage) leaves, and the suppressing effect was not as obvious as that at the four-leaf stage (Figure 1B and C). Protein P19 is a VSR encoded by tomato bushy stunt virus known for its systemic VSR activity. Compared with P19 and EV, P2 had systemic VSR activity (Figure 1D and E; Supporting Figure 2A and B); the systemic silencing efficiency of P2 exceeded 90% in each experiment repeat (Figure 1E; Supporting Figure 2B). These results demonstrated that accumulation level and VSR activity of P2 was lower in older than in younger *N. benthamiana* leaves, and P2 had systemic VSR activity in *N. benthamiana*.

**Figure 1.**
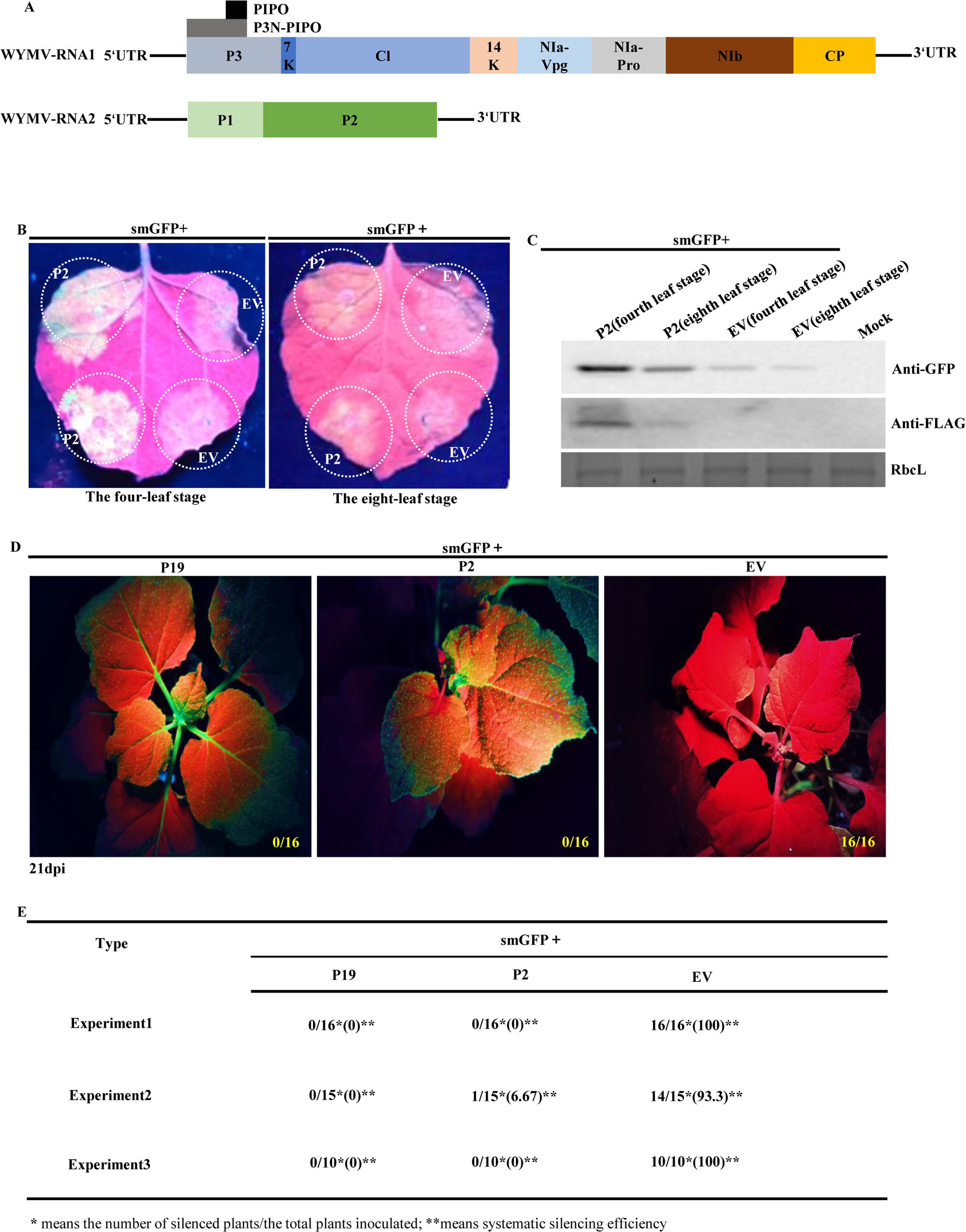
WYMV P2 protein has RNA-silencing suppressor activity. A, Genetic structure map of WYMV. B, WYMV P2 was cloned into an expression vector (pGD-3×FLAG or pMDC32-3×FLAG). The experimental group (P2+smGFP) and the control group (EV+smGFP) were infiltrated into *N. benthamiana* leaves at the eight- and four-leaf stages, respectively, in which EV was an empty vector; 3 days post-infiltration, they were observed under a UV lamp. C, Three days post-infiltration, the leaves of *N. benthamiana* in B were sampled, large subunit Rubisco (RbcL) of *N. benthamiana* was used as an internal control, and Anti-FLAG (rabbit) and Anti-GFP (rabbit) antibodies were used for western blotting. D, Positive control (P19+smGFP), experimental (P2+smGFP), and negative control (EV+smGFP) groups were respectively inoculated with equal numbers of *N. benthamiana* line 16c (three-leaf stage). The results were photographed under UV lamp at 21 days post-infiltration. E, Statistical data of the experimental results of three independent replicates of D.

To assess whether the P2 proteins of the remaining five viruses in *Bymovirus* (WSSMV, BaYMV, BaMMV, OMV, and RNMV; Supporting Figure 3A) have similar VSR activity, the WSSMV (NC_040507.1) and BaYMV (AJ132269.1) P2 genes were cloned. The GFP fluorescence intensity of WSSMV and BaYMV P2 proteins in *N. benthamiana* leaves at the four-leaf stage under a long-wave UV lamp was similar to that of WYMV P2 (Supporting Figure 3B), and these observations were confirmed by western blotting (Supporting Figure 3C). These results showed that WSSMV and BaYMV P2 proteins had strong local VSR activity on younger *N. benthamiana* leaves.

### Effects of WYMV P2 mutants on its VSR activity and its own protein stability

The WYMV P2 protein consists of 649 amino acids, and 10 truncated mutants were obtained successively: (AA1–222), (AA223–248), (AA249–649), (AA1–93), (AA94– 186), (AA187–279), (AA280–372), (AA373–465), (AA466–558), and (AA559–649). Of these, (AA1–222), (AA223–248), and (AA249–649) were constructed by predicting the protein structure on the InterPro website (http://www.ebi.ac.uk/interpro/), and the other mutants were obtained by truncating P2 with equal length (Figure 2A). Agrobacterium was infiltrated into *N. benthamiana* leaves at the four-leaf stage and 3 days later, the fluorescence intensity of the injection area containing the gene for P2 was similar to that of the injection area containing other mutants under a UV lamp (Figure 2B) – the western blotting results were consistent with these observations (Figure 2C). These results showed that P2 and its truncated mutants had strong local VSR activity on younger *N. benthamiana* (four-leaf stage) leaves.

**Figure 2.**
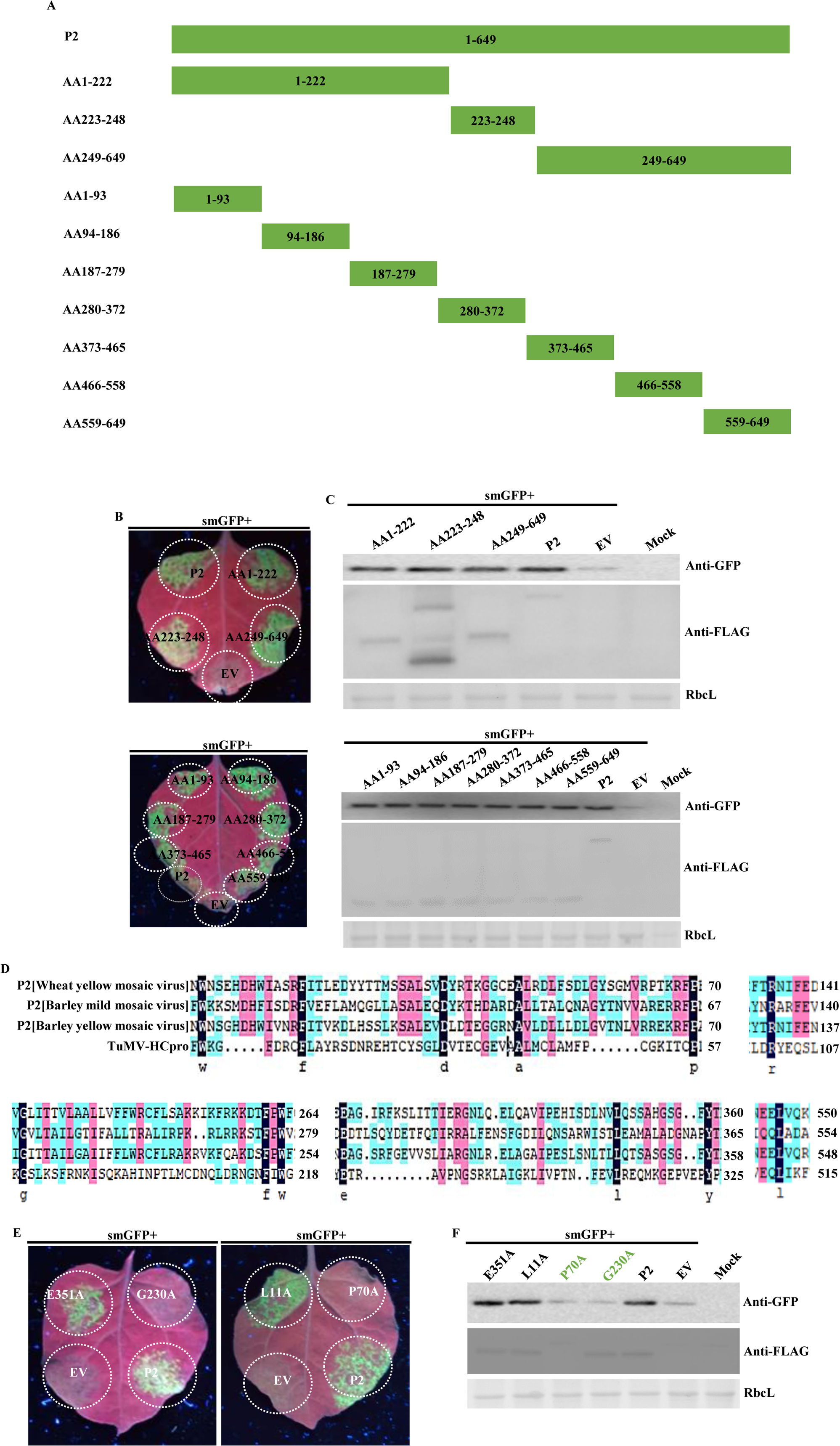
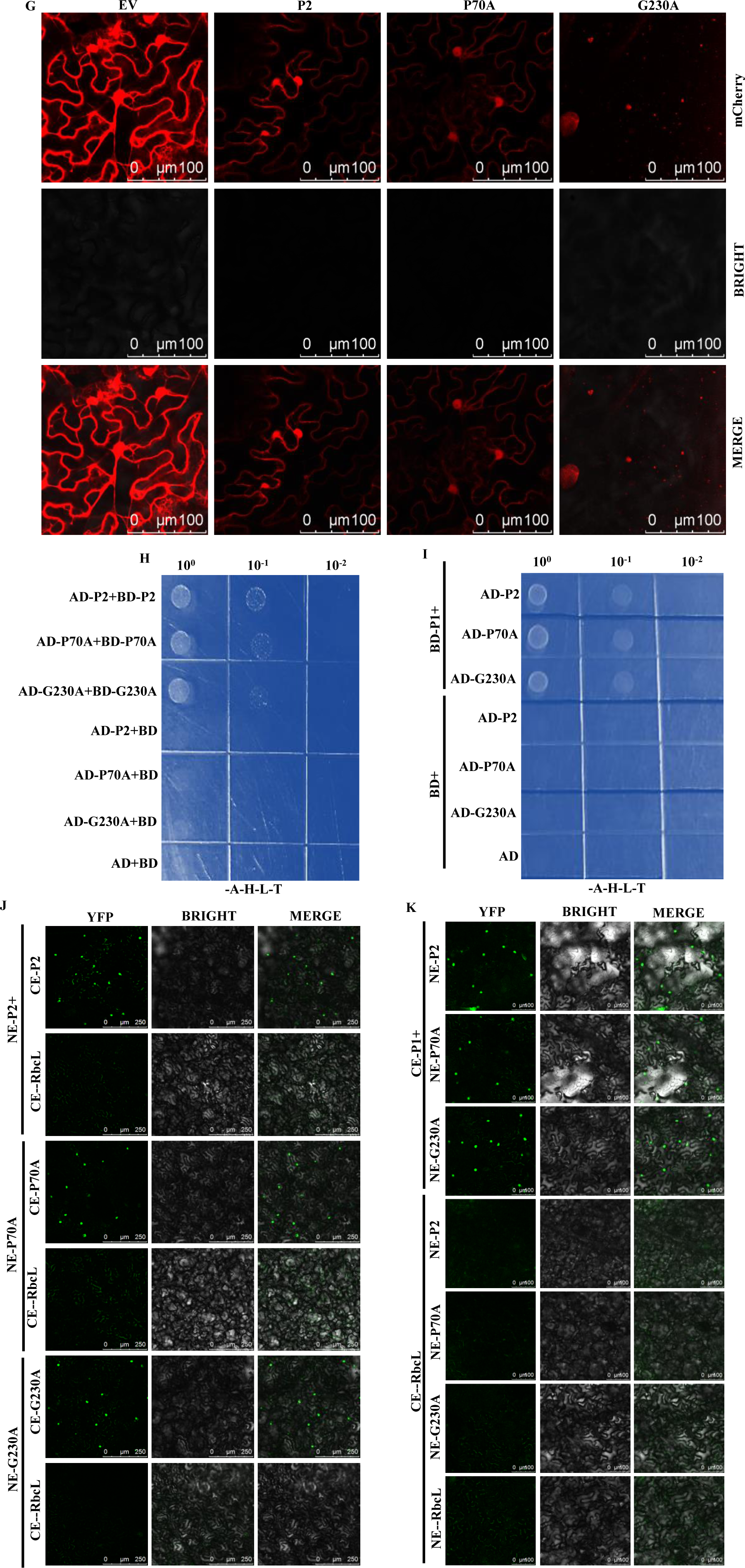
Effects of WYMV P2 mutants on its VSR activity and its own protein stability. A, Protein structure of P2 and its mutants; P2 and its mutants were tagged with 3×FLAG. B, Positive control (P2+smGFP), negative control (EV+smGFP), and experimental groups; all treatments were infiltrated into the same *N. benthamiana* leaf, and photographed under UV lamp 3 days post-infiltration. C, Three days post-infiltration, *N. benthamiana* leaves in B were sampled; large subunit RbcL of *N. benthamiana* was used as an internal control and Anti-FLAG (rabbit) and Anti-GFP (rabbit) antibodies were used for western blotting. D, Alanine scanning point mutations were introduced at 13 conserved amino acid sites. E, Thirteen point mutants of P2 in Agrobacterium suspensions were successively added with smGFP as 13 experimental groups, treatment (P2+smGFP) as a positive control group, and (EV+smGFP) as a negative control group; they were photographed under a UV lamp 3 days post-infiltration of leaves. F, Leaves containing mutants P70A and G230A were sampled, and then western blotting was performed with Anti-FLAG (rabbit) and Anti-GFP (rabbit) antibodies; the large subunit RbcL of *N. benthamiana* was used as an internal control. G, Subcellular localizations of P2, P70A, and G230A were observed. P2, P70A, and G230A were cloned into pGD-3G-mCherry. The P2-mCherry, P70A-mCherry, and G230A-mCherry were used as experimental groups, and pGD-3GmCherry was the control group; P19 was added to each group and then infiltrated into *N. benthamiana* leaves. H, Yeast two-hybrid (Y2H) experiment, treatment (AD+BD) was the negative control group; treatments (BD+AD-P2), (BD+AD-P70A), and (BD+AD-G230A) were the parallel control groups; and the rest were the experimental groups. The growth of yeast on deficient plates was recorded. I, Y2H experiment, treatment (AD+BD) was the negative control group; treatments (BD+AD-P2), (BD+AD-P70A), and (BD+AD-G230A) were the parallel control groups; and the rest were experimental groups. J, Bimolecular fluorescence complementation (BiFC) experiment, treatments (CE-P2+NE-P2), (CE-P70A+NE-P70A), and (CE-G230A+NE-G230A) were the experimental groups, and the rest were parallel control groups; P19 was added in each group. Samples were collected 3 days post-infiltration and observed under a confocal laser microscope. K: BiFC experiment, treatment (CE-RbcL+NE-RbcL) was the negative control group; treatments (CE-RbcL+NE-P2), (CE-RbcL+NE-P70A), and (CE-RbcL+NE-G230A) were the parallel control groups; and the rest were the experimental groups; P19 was added in each group.

Comparing amino acid sequences of P2 proteins from BaYMV, BaMMV, WYMY, and TuMV HC-Pro showed a total of 13 conserved amino acids (Figure 2D). Alanine point mutations were performed on the 13 conserved amino acids successively, and the loss of local VSR function of P70A and G230A was found by infiltration of the 13 point mutants (Figure 2E). Western blotting results showed that protein expression of G230A was consistent with that of P2, but expression of P70A was lower than that of P2 (Figure 2F). These results showed that P70A and G230A lost local suppressor function on younger *N. benthamiana* leaves, and the protein of P70A was more unstable.

Systemic VSR activity assays showed that G230A and mutants (AA1–222) and (AA249–649) lost systemic silencing suppressor activity (Supporting Figure 2A and B). Analysis of local and systemic VSR activities of P2, G230A, (AA1–222), and (AA249–649) showed that P2 had local and systemic VSR activities, but G230A had neither, and mutants (AA1–222) and (AA249–649) had local but no systemic VSR activity (Supporting Figure 2C).

There were obvious differences between the localization of P2 and G230A. Localization of G230A was more concentrated in the cytoplasmic aggregation body, while P2 and P70A were localized in both the nucleus and cytoplasm (Figure 2G). The yeast two-hybrid (Y2H) and bimolecular fluorescence complementation (BiFC) experiment showed that all the P2, P70A, and G230A could self-interact (Figure 2H and J), and interacted with P1 (Figure 2I and K).

Because we found that P70A protein was more unstable in *N. benthamiana*, the protein stability of P2 and its point mutants was investigated. Firstly, the protein accumulation of P2 in Dicer2,4RNAi *N. benthamiana* leaves was significantly higher than that in wild-type leaves (Supporting Figure 4A). When P2, P70A, and G230A were co-infiltrated with P19 into *N. benthamiana* leaves, protein accumulation of P2, P70A, and G230A was significantly higher than that without P19. Under the same conditions, protein accumulation of P70A was lower than that of P2, while G230A was consistent with P2 (Supporting Figure 4A), indicating that the RNA silencing of the host plants was not conducive to protein accumulation of P2 and its point mutants in *N. benthamiana* leaves. Secondly, western blotting results showed that protein accumulation of P2, P70A, and G230A increased at 8–16 °C compared with 26 °C (Supporting Figure 4B), indicating that low temperature was conducive to P2 accumulation in *N. benthamiana* leaves. Thirdly, the inhibitors 3MA, E64D, and MG132 significantly inhibited the degradation of P2 and G230A, and 3MA significantly inhibited the degradation of P70A (Supporting Figure 4C), indicating that autophagy and ubiquitination increased P2 protein accumulation in *N. benthamiana* leaves. Further experimental results showed that accumulation of protein P2 was significantly higher in ATG5-silenced (TRV:ATG5) than in GUS-silenced (TRV:GUS) *N. benthamiana* plants (Supporting Figure 4D), indicating that the autophagy indeed significantly upregulated P2 accumulation in *N. benthamiana*. These results showed that RNA silencing, high temperature, autophagy, and ubiquitination were all unfavorable to P2 accumulation in *N. benthamiana* leaves. Similarly, when WSSMV and BaYMV P2 proteins and P19 were co-injected into *N. benthamiana* leaves respectively, their protein accumulation was also upregulated compared with the control. When WSSMV and BaYMV P2 proteins were expressed at 8 ℃, their protein accumulation was higher compared with the 26 °C group. Inhibitor E64D significantly inhibited the degradation of WSSMV P2 and BaYMV P2 proteins, the accumulation of WSSMV P2 and BaYMV P2 proteins was increased compared with the control group (Supporting Figure 5A). Further results showed that TRV-silenced autophagy gene *ATG5* significantly upregulated accumulation of WSSMV and BaYMV P2 proteins in *N. benthamiana* leaves (Supporting Figure 5B). These results showed that RNA silencing, high temperature, and autophagy were not conducive to accumulation of WSSMV and BaYMV P2 proteins.

### WYMV P2 interacts with gene-silencing related protein NbCaM

We investigated possible mechanisms of WYMV P2 protein inhibiting RNA silencing. Three days after infiltration of *N. benthamiana* leaves with Agrobacterium suspension containing P2 vector (EV served as control group), the transcriptional expression levels of genes for key factors of the antiviral RNA-silencing pathway (e.g. CaM, CAMTA3, AGO1, SGS3, RDR6, and DCL2) were determined using quantitative real-time (qRT)-PCR. The results showed that WYMV P2 could reduce the expression levels of these key genes in the RNA-silencing pathway to varying degrees (Supporting Figure 6).

The qRT-PCR results showed that the gene for CaM was the most significantly downregulated; it was recently reported that VSRs exert suppressor activity by directly interacting with CaM, thereby interfering with the CaM–CAMTA3 interaction (Chen et al., 2023; Wang et al., 2021; Wang et al., 2022). Therefore, Y2H and BiFC were used to verify the P2–CaM interaction (Figure 3; Supporting Figure 7). The Y2H and BiFC experiments showed that P2 interacted with CaM, and the truncated mutants (AA1–222), (AA223–248), and (AA466–558) also interacted with CaM; however, P70A and G230A lost the ability to interact with CaM (Figure 3A and C). The Y2H and BiFC experiments showed that P2 could not interact with CAMTA3 (Figure 3B and D). Similarly, Y2H and BiFC experiments showed that P2 proteins of WSSMV and BaYMV also interacted with CaM, but not with CAMTA3 (Supporting Figure 8). These results demonstrated that P2 and its truncated mutants could interact with CaM, but not with CAMTA3, and the point mutants, P70A and G230A lost the ability to interact with CaM and CAMTA3.

**Figure 3.**
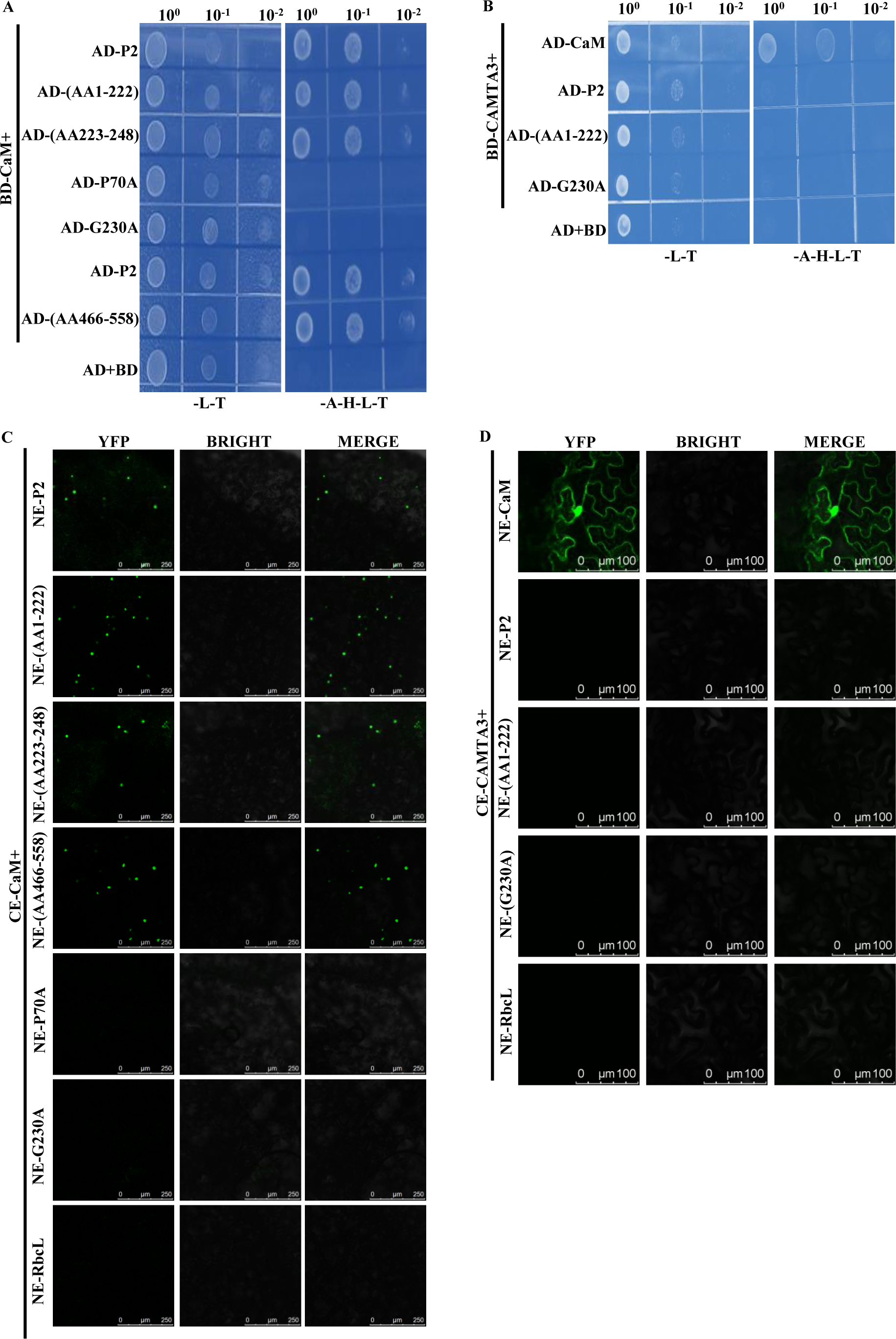
WYMV P2 interacts with the gene-silencing related protein NbCaM. A, Y2H experiment, treatment (AD+BD) was the negative control group, and the rest were the experimental groups. B, Y2H experiment, treatment (AD+BD) was the negative control group, and the rest were the experimental group. C, BiFC experiment; treatment (CE-CaM+NE-RbcL) was the parallel control group, and the rest were experimental groups; P19 was added to each group. Samples were collected 3 days post-infiltration and observed under a confocal laser microscope. D, BiFC experiment; treatment (CE-CAMTA3+NE-RbcL) was used as the parallel control group, and the rest were experimental groups; P19 was added to each group. Samples were collected 3 days post-infiltration and observed under a confocal laser microscope.

### WYMV P2 interferes with the interaction of gene-silencing related proteins NbCaM and NbCAMTA3

Since WYMV P2 can interact with CaM, it is necessary to verify whether P2 can also interfere with the CaM–CAMTA3 interaction. The results of competitive BiFC and competitive co-immunoprecipitation (Co-IP) assays showed that P2, and truncated mutants (AA223–248) and (AA1–222), could significantly interfere with the CaM– CAMTA3 interaction. Point mutant G230A lost the ability to interfere with the CaM– CAMTA3 interaction (Figure 4A and B). In addition, WSSMV and BaYMV P2 proteins also interfered with the CaM–CAMTA3 interaction (Supporting Figure 9). These results demonstrated that P2, and truncated mutants (AA223–248) and (AA1– 222), could interfere with the CaM–CAMTA3 interaction to inhibit RNA silencing, while G230A could not interfere with the interaction.

**Figure 4.**
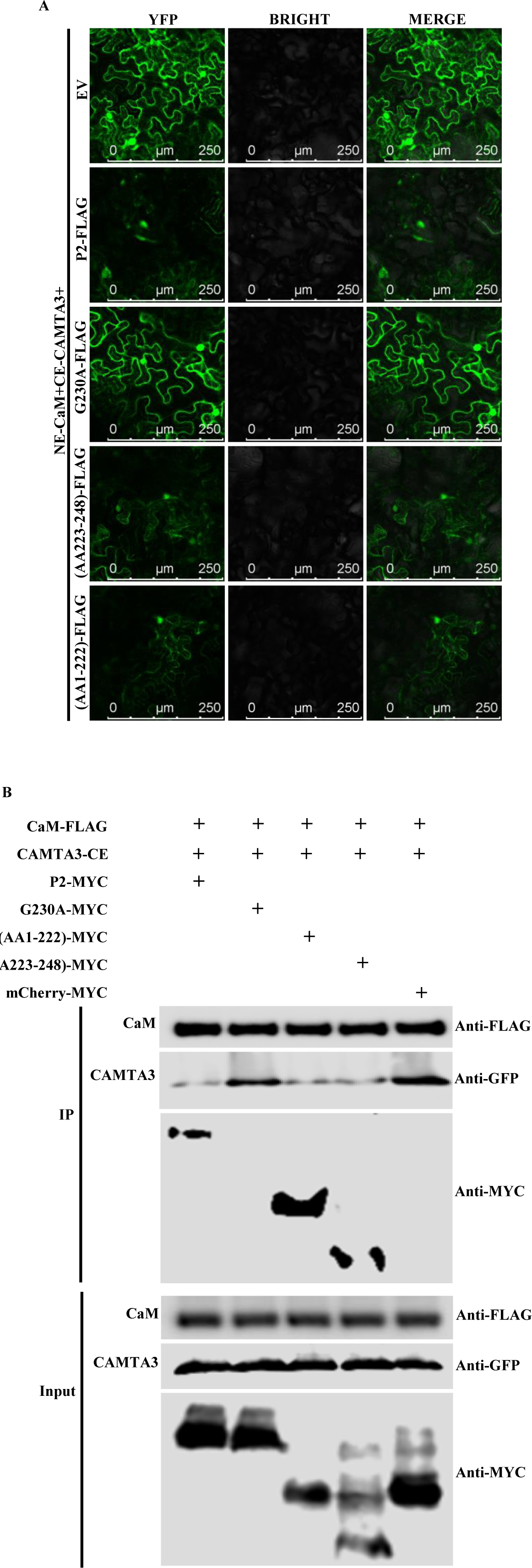
WYMV P2 interferes with the interaction of gene-silencing related proteins NbCaM and NbCAMTA3. A, Competitive BiFC analysis of P2 interference with CaM–CAMTA3 interaction; treatment (EV+NE-CaM+CE-CAMTA3) as the control group, and the rest as experimental groups. Samples were collected 3 days post-infiltration and observed under a laser confocal microscope. B, Competitive co-immunoprecipitation (Co-IP) analysis of P2 interfering with CaM–CAMTA3 interaction; treatment (mCherry-MYC+CAM-FLAG+CE-CAMTA3) was the control group, and the rest were experimental groups. Samples were collected 3 days post-infiltration and Co-IP experiments were conducted with FLAG beads.

In addition, we observed the colocalization of CaM respectively with P2, P70A, and G230A. The P2 and CaM were co-located in cytoplasm and nucleus, P70A and CaM were co-located in cytoplasm and nucleus, while G230A and CaM were mainly co-localized in cytoplasmic aggregation bodies (Supporting Figure 10).

### WYMV P2 VSR activity participates in virus infection of wheat

To determine whether the VSR-deficient mutants of P2 affect WYMV infection, directed mutation sites were generated in the infectious clone pWY2 (RNA2 of WYMV) of the Japanese isolate, and the mutants pWY2 (P70A) and pWY2 (G230A) were obtained (Figure 5A). Transcription bands of pWY1 (RNA1 of WYMV), pWY2, pWY2 (P70A), and pWY2 (G230A) were detected using agarose gel electrophoresis (Figure 5B). The pWY1 transcript was abbreviated as RNA1, pWY2 transcript as RNA2, pWY2 (P70A) transcript as RNA2 (P70A), and pWY2 (G230A) transcript as RNA2 (G230A). The upper leaves of wheat plants inoculated with wild-type transcripts mixture (RNA1+RNA2) showed severe yellowing; the upper leaves of wheat plants inoculated with the WYMV-P70A mutant transcripts mixture (RNA1+RNA2[P70A]) and those inoculated with the WYMV-G230A mutant transcripts mixture (RNA1+RNA2[G230A]) showed mild yellowing (Figure 5C). The upper non-inoculated leaves were collected for semi-quantitative RT-PCR, which showed that the accumulation of WYMV RNA1 and RNA2 of the wild type was higher than that of WYMV-P70A and WYMV-G230A in both positive and negative chains. Western blotting results showed that the CP protein accumulation of the wild type was higher than that of WYMV-P70A and WYMV-G230A, and the P2 protein accumulation of WYMV-G230A did not significantly differ from that of the wild type, while P2 protein accumulation of WYMV-P70A was the lowest (Figure 5D). The RT-PCR product sequencing confirmed that the progeny virus RNA2 and its mutants retained their original nucleotide sequence. The results of *in vitro* transcript inoculation of WYMV infectious clone showed that P2 VSR activity played a key role in WYMV infection, VSR defective mutants P70A and G230A reduced WYMV infection in wheat, and P70A was still relatively unstable in wheat plants as it was in *N. benthamiana* (Figure 5).

**Figure 5.**
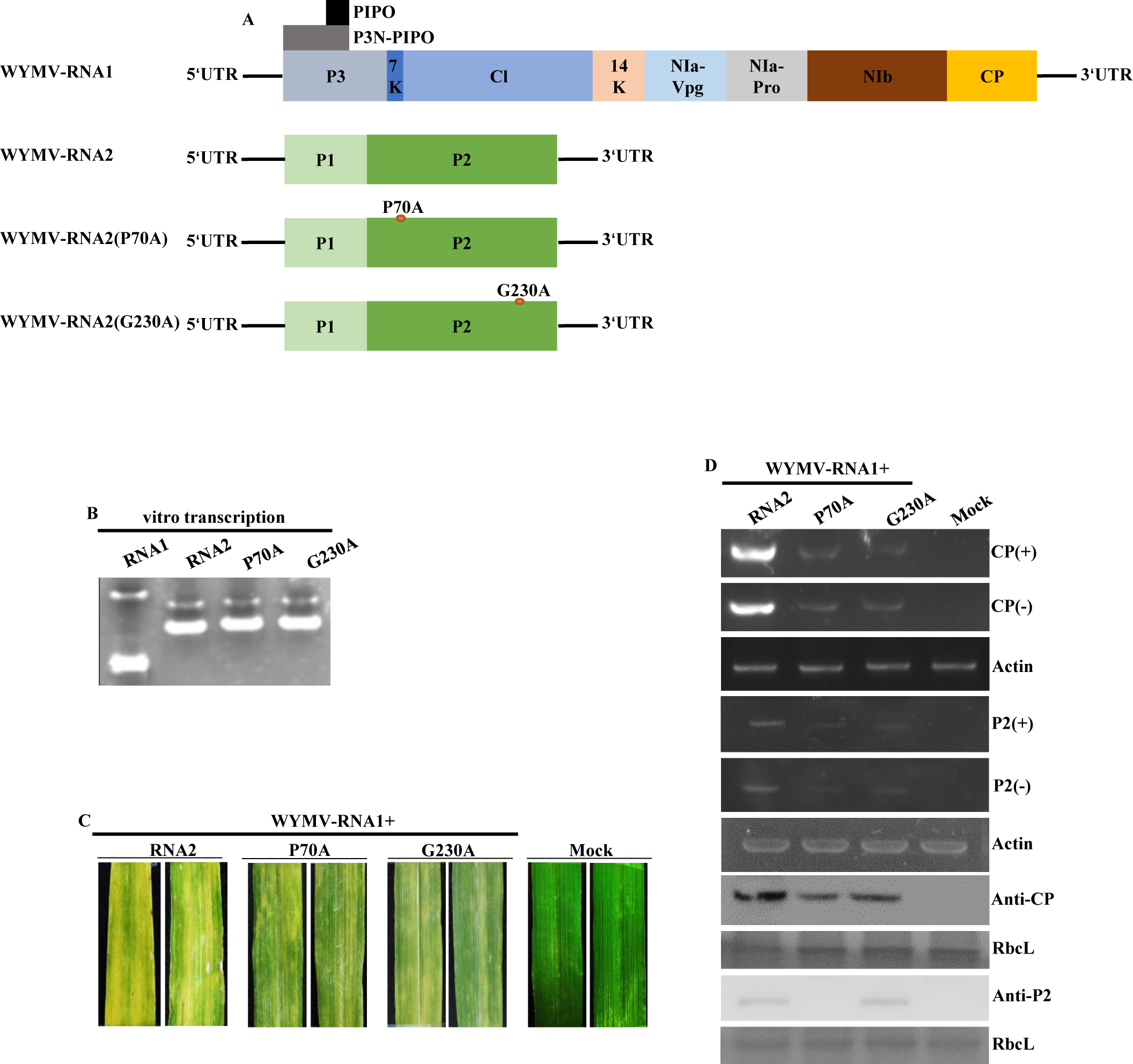
WYMV P2 VSR activity is involved in virus infection of wheat. A, Structure diagram of the infectious clone WYMV RNA1 (pWY1) and RNA2 (pWY2) under the control of the SP6 promoter, and two point mutants P70A and G230A constructed based on RNA2 (pWY2). B, *In vitro* transcription of pWY1, pWY2, pWY2 (P70A), and pWY2 (G230A). First, pWY1, pWY2, pWY2 (P70A), and pWY2 (G230A) were linearized with *Spe*I enzyme, and then transcribed *in vitro*. The transcription reaction was performed at 37 ℃ for 1 h, followed by the addition of 1 μL of enzyme for another 1 h, and the transcripts were detected by agarose gel electrophoresis. C, The pWY1 transcript (hereinafter referred to as RNA1) was respectively mixed with the pWY2, pWY2 (P70A), and pWY2 (G230A) transcripts (referred to as RNA2, RNA2[P70A], and RNA2[G230A], respectively) to ensure that the concentrations of each transcript were equal in different groups. Then an equal volume of 2×FES buffer was added to each treatment for mixing, including wild-type WYMV (RNA1+RNA2+FES buffer), WYMV-P70A (RNA1+RNA2[P70A]+FES buffer), WYMV-G230A (RNA1+RNA2[G230A]+FES buffer), and mock inoculation group (mock, FES buffer). Six wheat seedlings were inoculated in each experimental group. After mechanical inoculation, wheat plants were cultured at 8 ℃ for 15 days, then at 25 ℃ for 15 days, and then the phenotype was photographed. D, Upper non-inoculated leaves were sampled, and semi-quantitative RT-PCR and western blotting were used for detection. Wheat *Actin* was used as an internal reference for RT-PCR detection, and wheat large subunit RbcL was used as an internal reference for western blotting.

## Discussion

The results demonstrated that WYMV P2 had local and systemic VSR activities. Agrobacterium co-infiltration experiments showed that P2 had relatively strong local VSR activity in *N. benthamiana* leaves at the four-leaf stage, but weak at the eight-leaf stage (Figure 1B and C). The WYMV P2 was similar to P1 in that it had similar local VSR activity and only exhibited strong suppressor activity in the younger *N. benthamiana* leaves, which further explains why wheat seedlings need to be inoculated with WYMV at the one to two leaf stage (Chen et al., 2023; Zhang et al., 2021; Shang et al., 2002). One significant difference between them was that the P2 truncated mutants had local VSR activity (Figure 2A–C), while the P1 truncated mutants tested lost local VSR activity (Chen et al., 2023). Amino acid sequence alignment of *Bymovirus* P2 revealed 13 conserved amino acid residues (Figure 2D), and VSR activity screening of the 13 point mutants showed that conserved proline 70 and glycine 230 in P2 were essential for RNA-silencing suppressor activity (Figure 2E and F). Point mutants P70A and G230A lost local VSR activity, but the Y2H and BiFC experiments showed that they maintained protein self-interaction, indicating that the protein self-interaction may not be sufficient for local VSR activity (Figure 2H and J). Interestingly, the Y2H and BiFC experiments showed that P70A and G230A maintained interaction with P1 protein, suggesting that the loss of P2 local VSR activity did not depend on its interaction with P1 protein (Figure 2I and K).

Viruses have developed a variety of mechanisms to overcome antiviral RNA silencing. In general, the mechanisms of viral silencing suppressors can be divided into two broad categories in the downstream of the RNA-silencing pathway (Brodersen and Voinnet, 2006; Chen et al., 2023; Li et al., 2019; Marco et al., 2013). The first kind of VSRs act on proteins associated with RNA silencing, such as the P38 protein of turnip crinkle virus (TCV) (Schott et al., 2012; Thomas et al., 2003), the P1 protein of sweet potato mild mottle virus (Giner et al., 2010) interacts with AGO1 to inhibit RNA silencing, V2 encoded by tomato yellow leaf curl virus interacts with SGS3 and exerts suppressor activity (Glick et al., 2008), and P6 protein encoded by rice yellow stunt virus interacts with RDR6 to inhibit transgene-induced systemic silencing (Hongyan et al., 2013). The second kind is VSRs that act on RNA via the RNA-silencing pathway, such as P14 protein encoded by pothos latent virus and the P38 protein encoded by TCV and B2 protein encoded by flock house virus, which can bind to dsRNA (Chao et al., 2005; Merai et al., 2005; Merai et al., 2006). It has been reported that VSR protein negatively regulates host RNA silencing by binding to CaM (Nakamura et al., 2014). Recent research results further revealed that some VSRs can play a role in the upstream of RNA-silencing pathway. Wang et al. (2021) found that VSR protein V2 encoded by cotton leaf curl multan virus (CLCuMuV) can promote viral replication by disrupting the NbCaM–NbCAMTA3 interaction, and V2 can competitively bind CaM and inhibit the Ca^2+^–CaM–CAMTA3 signaling cascade, thereby inhibiting the antiviral response and exerting its suppressor activity. Our laboratory also found that WYMV P1 can interact not only with CaM, but also with CAMTA3, thereby interfering with the CaM–CAMTA3 interaction to exert VSR activity (Chen et al., 2023). In the present study, WYMV P2 interacted with CaM protein (Figure 3A and C) to interfere with the CaM–CAMTA3 interaction (Figure 4A and B), while P70A and G230A did not interact with CaM and did not to interfere with the CaM–CAMTA3 interaction, suggesting that CaM may play a key role in the silencing of antiviral RNA infected by WYMV. Interestingly, WYMV P1 can interact with CaM and CAMTA3 (Chen et al., 2023); however, P2, like CLCuMuV V2, can interact with CaM only, not with CAMTA3 (Figure 3; Wang et al., 2021). All the P2 truncated mutants tested showed local VSR activity (Figure 2A–C), and the truncated mutants of (AA1–222) and (AA223–248) interacted with CaM to interfere with the CaM–CAMTA3 interaction (Figures 3 and 4A and B), while the P1 truncated mutant (AA16–255) lost the ability to interfere with the CaM–CAMTA3 interaction (Chen et al., 2023), which partly explains the uniqueness of P2 as opposed to P1.

Sun et al. (2013) reported that eGFP-P2 was localized in both cytoplasm and nucleus. We noted that point mutant G230A formed cytoplasmic aggregation bodies in the cytoplasm, differing from the localization of P2 and P70A in the cytoplasm and nucleus (Figure 2G). The P2 and P70A co-located with CaM in the cytoplasm and nucleus, respectively, whereas G230A co-located with CaM mainly in cytoplasmic aggregation bodies (Supporting Figure 10). The results of intracellular localization experiments suggest that P2 and CaM colocalization in the nucleus may be related to VSR activity.

In addition, accumulation of WYMV P2 protein in *N. benthamiana* leaves was significantly affected by temperature, with the low temperature range of 8–16 °C conducive to P2 accumulation in leaves (Supporting Figure 4B). Relevant research showed that WYMV systemically infected wheat seedlings at 8 ℃ (Zhang et al., 2021), and Japanese scholars reported that the inoculation conditions of WYMV and BaYMV *in vitro* transcripts were 8–12 and 15 ℃, respectively (Ohki et al., 2019; You and Shirako, 2010). These results are consistent with the 8–15 ℃ required for mechanical inoculation of WYMV (Shang et al., 2002; Yang et al., 2022). Related articles reported that low temperature can reduce the function of gene silencing in plants (Fei et al., 2021; Sos-Hegedus et al., 2005; Tuttle et al., 2008), which are similar to WYMV P1, with low temperature conducive to accumulation of P2 protein (Supporting Figure 4B; Chen et al., 2023) – this partly explains why suitable low-temperature conditions are required to establish WYMV infection. Both P2 and G230A had similar accumulation levels at low and normal temperatures (Supporting Figure 4B), indicating that point mutations in G230A did not affect their protein accumulation levels. We also found that P2 protein accumulation in *N. benthamiana* leaves was significantly reduced through autophagy but, unlike P1, P2 can also be degraded by ubiquitination (Supporting Figure 4C; Chen et al., 2023). Autophagic inhibitors had a significant effect on P2 and G230A accumulation, suggesting that this point mutation did not affect P2 degradation by autophagy (Supporting Figure 4C). 3MA is a classic autophagy inhibitor, which mainly plays an inhibitory role in the formation and development of autophagosomes (Seglen and Gordon, 1982; Xiumei et al., 2019). Interestingly, P70A protein itself is relatively unstable, and only 3MA inhibits P70A degradation (Supporting Figure 4C), which may make P2 more susceptible to autophagy at the vesicle nucleation stage. The TRV system silenced the autophagy gene *ATG5* in *N. benthamiana* leaves, significantly upregulating accumulation of P2 protein in leaves (Supporting Figure 4D). Silencing of ATG5 leads to the degradation of autophagy hexamers containing ATG12 complex, and autophagosomes only form normally with difficulty (Yang et al., 2018). Therefore, compared with control plants, accumulation of P2 in ATG5-silenced *N. benthamiana* plants was significantly upregulated (Supporting Figure 4D). Compared to WYMV P1, P2 protein in *N. benthamiana* is degraded by not only autophagy but also ubiquitination pathways, which may partially explain why a WYMV infectious clone cannot systemically infect *N. benthamiana* (Chen et al., 2023; Zhang et al., 2021), and why WYMV can only systemically infect wheat under natural conditions (Clover and Henry, 2004; Li and Shirako, 2015).

In summary, we proposed a VSR working model of WYMV P2 (Figure 6). In this model, WYMV P2 protein exerts VSR activity by interfering with the CAM–CAMTA3 interaction in host plants. Among them, P2 protein exerts VSR activity by interfering with the CaM–CAMTA3 interaction, and promotes systemic infection of WYMV in wheat. Neither P70A nor G230A could interact with CaM, and P70A and G230A lost their ability to interfere with the CaM–CAMTA3 interaction, ultimately resulting in attenuating WYMV infection in VSR-deficient wheat mutants P70A and G230A. These results provide clues for further elucidating the underlying molecular mechanism of WYMV interaction with wheat and effective prevention and management of the diseases caused by this virus.

**Figure 6.**
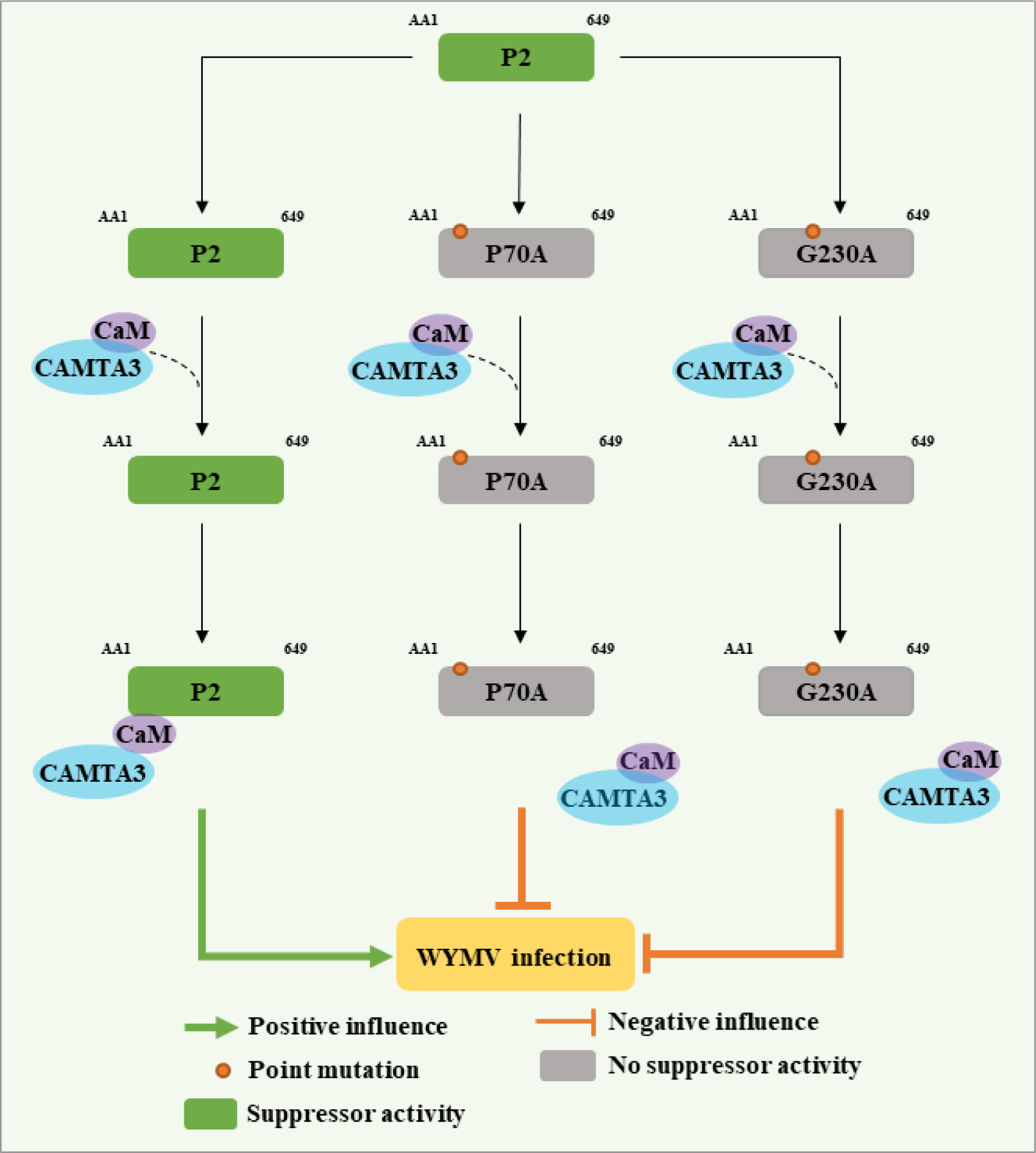
Working model showing that VSR activity of WYMV P2 promotes WYMV infection. WYMV P2 protein displays VSR activity by interfering with CaM– CAMTA3 interaction in host plants, and promotes WYMV to systemically infect wheat plants. Point mutants P70A and G230A cannot interact with CaM and cannot interfere with CaM–CAMTA3 interaction, resulting in the loss of VSR activity of both P70A and G230A, which significantly reduces the systemic infection efficiency of WYMV. In summary, P2 plays a VSR role in wheat and promotes WYMV infection by interfering with CaM-related antiviral RNAi defense pathway.

## Materials and methods

### Plant materials and growing conditions

Transgenic *N. benthamiana* with GFP gene (line 16c), DCL2,4-RNAi *N. benthamiana*, and wild-type *N. benthamiana* were grown in a greenhouse with a light cycle of light/dark of 16/8 h, temperature of 25 ℃, and relative humidity of 60% (Li et al., 2019; Zhuo et al., 2014). After inoculation with transcripts of the WYMV infectious clone, wheat (variety Yangmai 158) seedlings at one-leaf stage were first cultured at 8 ℃ for 15 days and then at 25 ℃ for 15 days with a constant relative humidity of 60% (Chen et al., 2023).

### Chemical reagents and enzymes

Autophagic inhibitors 3MA and ED64 were provided by Selleck Chemicals (Houston, TX, USA). Ubiquitination inhibitor MG132 was purchased from Merck Calbiochem (Billerica, MA, USA). The plasmid extraction kit was purchased from Axygen (Hangzhou, China). Gel recovery kit was purchased from Omega (Guangzhou, China). *Escherichia coli* culture-related reagents and yeast were purchased from Biodee (Beijing, China). TRIzol reagent was purchased from Invitrogen (Shanghai, China). Protein marker was purchased from Fermentas (Beijing) and M-MLV reverse transcriptase was purchased from Promega (Beijing). Homologous recombinant kit was purchased from Clone Smarter (Beijing). Agarose and DNA marker were purchased from Fermentas (Shanghai). The 2×M5 HiPer Realtime PCR Super mix was purchased from Mei5 Biotechnology (Beijing). Primer Star high fidelity DNA polymerase and restriction enzymes were purchased from TaKaRa (Beijing). Ordinary Taq enzyme was purchased from Tiangen (Beijing).

### Strains and vectors

The WYMV infectious clones under control of the SP6 promoter are referred to as pWY1 and pWY2 (Chen et al., 2023; Li and Shirako, 2015). The two *P2* genes [WSSMV (*P2*) and BaYMV (*P2*)] and *NbCaM* gene were synthesized by Shanghai Gerry Biological Company. *Agrobacterium tumefaciens* strains EHA105 and GV3101 were purchased from Beijing DIA-UP Biotechnology Company. *Escherichia coli* MC1022 was a gift from Dr. Salah Bouzoubaa (Li et al., 2019). The plasmids TRV:GUS and TRV:NbATG5 were produced in a research laboratory (Li et al., 2019). Plasmid NbCAMTA3 was provided by Professor Liu Yule (Wang et al., 2021). The BiFC vectors pSPYNE (NE) and pSPYCE (CE) were presented by Professor Joerg Kudla of Germany (Waadt et al., 2008; Walter et al., 2004). The Y2H vectors pGADT7 (AD) and pGBKT7 (BD) were purchased from Clontech (Beijing). The pGD-3GmCherry was a vector modified by pGD (Sun et al., 2018). The pMDC32-3×FLAG was produced in a research laboratory (Curtis and Grossniklaus, 2003). The pGD series vectors and pGDP19 were provided by Professor Andrew O. Jackson (Goodin et al., 2002).

### Antibodies

The WYMV CP and P2 antisera were prepared in our laboratory. The FLAG monoclonal antibody was purchased from Sigma (Saint Louis, MO, USA), and anti-rabbit secondary antibodies coupled with alkaline phosphatase or horseradish peroxidase were purchased from Sigma. The GFP and mCherry polyclonal antibodies were purchased from Bioeasytech (Beijing).

### Primers synthesis and sequencing

Primers were synthesized and sequenced by Ruibiotech (Beijing) and Sinogeno (Beijing).

### Point mutation and homologous recombination

The remaining plasmids were constructed by homologous recombination kits or point mutation kits. The homologous recombination experiment was performed with the Clone Smarter Seamless Connection Kit, and the specific experimental operation was according to Chen et al. (2023). The plasmid construction of point mutation was conducted using a Tiangen kit, and the primer design was with a Tiangen kit (https://www.docin.com/p-1670442438.html). All primers used are listed in Supporting Table 1.

### Agrobacterium transformation and infiltration

The optical density (OD) value of the sample was 0.5, while the OD value of P19 was 0.2. The sample was left for 2 h, and then infiltrated into *N. benthamiana* leaves. For specific operation methods, refer to Zhang et al. (2017).

### Total RNA extraction, RT-PCR, and qRT-PCR detections

Total RNA from plants was extracted with TRIzol reagent from Invitrogen (Carlsbad, CA, USA), and then RT-PCR or qRT-PCR was performed. For specific methods, refer to Jiang et al. (2020).

### Observation and photography under UV light

A digital camera (Canon Company, Tokyo, Japan) was used to photograph GFP fluorescence on *N. benthamiana* leaves under a UV lamp (B-100AP/ R; UVP Inc., Upland, CA, USA). For specific operation methods, refer to Zhuo et al. (2014).

### Inhibitors treatment experiment

The treatment methods of inhibitors (MG132, 3MA, and E64D) were tested according to Chen et al. (2023) and Li et al. (2019).

### Virus-induced gene silencing (VIGS) assay

For detailed VIGS experiment procedures, refer to Li et al. (2019) and Liu et al. (2002).

### Western blotting

The western blotting procedure was according to Sun et al. (2018).

### Observation by confocal microscope

The microscope used was a Leica SP8 (Leica, Frankfurt, Germany). The excitation wavelength of mCherry was 546 nm, and that of GFP was 488 nm. For detailed operation methods, refer to Jiang et al. (2020).

### BiFC and competitive BiFC experiments

The excitation wavelength of YFP was 514 nm, and the specific experimental methods of BiFC and competitive BiFC were according to Chen et al. (2023), Jiang et al. (2020), and Wang et al. (2021).

### Y2H experiment

The specific experimental method of Y2H followed Chen et al. (2023) and Zhang et al. (2017).

### Competitive Co-IP assay

For the specific procedure of competitive Co-IP experiment, refer to Chen et al. (2023) and Yang et al. (2018).

### *In vitro* transcription and inoculation

For specific methods of *in vitro* transcription and inoculation of wheat, refer to Chen et al. (2023) and Li and Shirako (2015).

## Authors’ contributions

C-G Han and D Chen designed the experiments. D Chen completed the experiments with the help of H-Y Zhang. S-M Hu, Z He, Y-Q Wu, Z-Y Zhang, Y Wang, and C-G Han analyzed the data and contributed through discussions; D Chen and C-G Han wrote the paper. All authors read and agreed to the publication of the manuscript.

## Acknowledgments

This research was supported by the National Science and Technology Research Project (2016ZX08002-001). We thank Dr. David C. Baulcombe (University of Cambridge, UK) for providing *N. benthamiana* and GFP transgenic *N. benthamiana* 16c; Dr. Andrew O. Jackson (University of California, USA) for providing pGDP19, pGDG, and pGD vectors; Dr. Salah Bouzoubaa for giving *E. coli* MC1022; and Prof. Yule Liu for providing NbCAMTA3 plasmid. Thanks to Professor Li-ying Sun for the gift of WYMV infectious clone. Thanks to Professors Xian-bing Wang, Yong-liang Zhang, Da-wei Li, Jia-lin Yu, and Sek-Man Wong for their guidance on this topic.

## Data availability

All relevant data can be found within the manuscript and its supporting materials.

## Supporting Information

**Supporting Figure 1** WYMV P1 and P2 sequences were compared with TuMV HC-Pro respectively.

**Supporting Figure 2** WYMV P2 has systemic RNA-silencing suppressor activity.

**Supporting Figure 3** WSSMV and BaYMV P2 proteins also exhibit local VSR activity.

**Supporting Figure 4** RNA silencing, temperature, and autophagy have significant effects on WYMV P2 protein accumulation.

**Supporting Figure 5** RNA silencing, temperature, and autophagy have significant effects on accumulation of WSSMV and BaYMV P2 proteins.

**Supporting Figure 6** WYMV P2 protein can reduce mRNA expression levels of RNA-silencing related genes such as those for CaM and CAMTA3 in *N. benthamiana* leaves to varying degrees.

**Supporting Figure 7** Preliminary experiments showed that WYMV P2 interacts with gene-silencing associated protein NbCaM.

**Supporting Figure 8** WSSMV and BaYMV P2 proteins can also interact with the gene-silencing associated protein NbCaM.

**Supporting Figure 9** WSSMV and BaYMV P2 proteins can also interfere with the NbCaM–NbCAMTA3 interaction.

**Supporting Figure 10** Colocalization of WYMV P2 and its mutants repectively with NbCaM.

**Supporting Table 1** Primers used in this manuscript.

